# Revealing Molecular Mechanisms of Early-Onset Tongue Cancer by Spatial Transcriptomics

**DOI:** 10.1101/2023.10.12.562054

**Authors:** Marina R. Patysheva, Elena S. Kolegova, Anna A. Khozyainova, Elizaveta A. Prostakishina, Maxim E. Menyailo, Irina V. Larionova, Oleg I. Kovalev, Marina V. Zavyalova, Irina K. Fedorova, Denis E. Kulbakin, Andrey P. Polyakov, Liliya P. Yakovleva, Mikhail A. Kropotov, Natalya S. Sukortseva, Yusheng Lu, Lee Jia, Rohit Arora, Evgeny L. Choinzonov, Pinaki Bose, Evgeny V. Denisov

## Abstract

Tongue cancer at a young age demonstrates an increase in incidence, aggressiveness, and poor response to therapy. Classic etiological factors for head and neck tumors such as tobacco, alcohol, and human papillomavirus are not related to early-onset tongue cancer. Mechanisms of development and progression of this cancer remain unclear. In this study, we performed spatial whole-transcriptome profiling of tongue cancer in young adults compared with elderly patients. Oxidative stress, vascular mimicry, and MAPK and JAK-STAT pathways were enriched in early-onset tongue cancer. Tumor microenvironment demonstrated increased gene signatures corresponding to myeloid-derived suppressor cells, tumor-associated macrophages, and plasma cells. The invasive front was accompanied by vascular mimicry with arrangement of tumor-associated macrophages and aggregations of plasma cells and lymphocytes organized into tertiary lymphoid structures. Taken together, these results indicate that early-onset tongue cancer has distinct spatial transcriptomic features and molecular mechanisms compared to older patients.

**HIGHLIGHTS:**

- Early-onset tongue cancer demonstrates extremely downregulated oxidative phosphorylation and upregulated glycolysis.
- MAPK pathway is the key player in the pathogenesis of tongue cancer in young adults.
- Early-onset tongue cancer is characterized by JAK-STAT dependent vascular mimicry supported by tumor-associated macrophages at the invasive edge.
- Tongue cancer microenvironment in young adults enriches for immunosuppressive myeloid derived suppressor cells and demonstrates reduced antigen presentation function.
- The tumor border in early-onset tongue cancer is enriched with plasma cells and lymphocytes in tertiary lymphoid structures.

## INTRODUCTION

Squamous cell carcinoma of the oral cavity has historically been considered a disease more common in individuals in their sixth decade of life that is related to cumulative exposure to tobacco, alcohol, and human papillomavirus (HPV) infection [1]. However, the first cases of this cancer in young adults were described in the 1930s, and the incidence of early-onset tongue squamous cell carcinoma (TSCC), especially in women, is increasing every year [2]. The etiological factors and mechanisms of TSCC initiation and progression in young adults have been the subject of significant debate and controversy owing to a lack of association with tobacco, alcohol consumption, and HPV [3].

Single studies showed that some cases with early-onset TSCC were related to the KIR2DL1^+^-HLA-C2^+^ genotype and MMP-1 2G allele. Molecular profile of early-onset TSCC are characterized by alterations in the *TP53, CDKN2A, CASP8, NOTCH1, FAT, ATXN1*, and *CDC42EP1* genes and methylation changes in the *RASSF1A, RASSF2A, MGMT, DAPK*, and *FHIT* genes [3-7]. Bulk RNA sequencing demonstrated upregulation of the immunosuppressive *LAG-3* and *HAVCR2* genes in early-onset TSCC [8]. By analyzing the The Cancer Genome Atlas (TCGA) data, we found that TSCC in young adults displays reduced mutational burden, overexpression of the Rap1, PI3K-Akt, MAPK, JAK-STAT, cGMP-PKG and Fc-gamma R-mediated phagocytosis signaling genes, and specific microbiome [9]. Nevertheless, a comprehensive picture of the pathogenesis of TSCC in young adults is still unclear. This can be due to a lack of investigations of tumor tissue as a complex system considering both tumor and microenvironment cell populations and their interactions.

Spatial transcriptomics technologies make it possible to analyze the transcriptome of individual cells with their original localization within the tissue. A recent study developed a spatial transcriptome map of early-stage oral squamous cell carcinoma, adjacent precancerous lesions, and a matched normal region [10]. Spatial transcriptomics also revealed differences in the biology of core and edge regions of oral tumors and identified patterns of expressed genes associated with disease prognosis [11].

In this work, we set out to investigate spatial transcriptomics of early-onset TSCC relative to this cancer in older adults. We uncovered a set of MAPK and JAK-STAT signaling pathways genes in the tumor cells and a decrease in antigen presentation, alongside myeloid derived suppression cells and tumor-associated macrophages with their synergistic efforts promoting vascular mimicry. Plasma cells and lymphocytes in tertiary lymphoid structures were enriched in the invasive front of TSCC in young adults. Together, our results highlight potential mechanisms of early-onset TSCC and serve as a reference for further studies of this cancer.

## RESULTS

### Spatial transcriptome profiling of early-onset TSCC

Tumor samples from 5 young and 4 elderly patients were used to investigate how early-onset TSCC differs from late-onset ones in spatial transcriptomics (Figure 1A). First at all, we performed the integration of spatial transcriptomics data from FFPE and FF samples. It is worth noting that the data included in the study differed significantly in the method of transcript capture. Molecular profiling of FF samples was carried out based on total mRNA, while profiling of FFPE samples was performed using probes targeting protein coding genes. To solve this problem, we removed genes from FF samples that were not captured by FFPE probes prior to integration. Further use of the Harmony tool gave us the opportunity to perform integration with sufficient quality for further analysis (Figure 1A).

**Figure 1.**
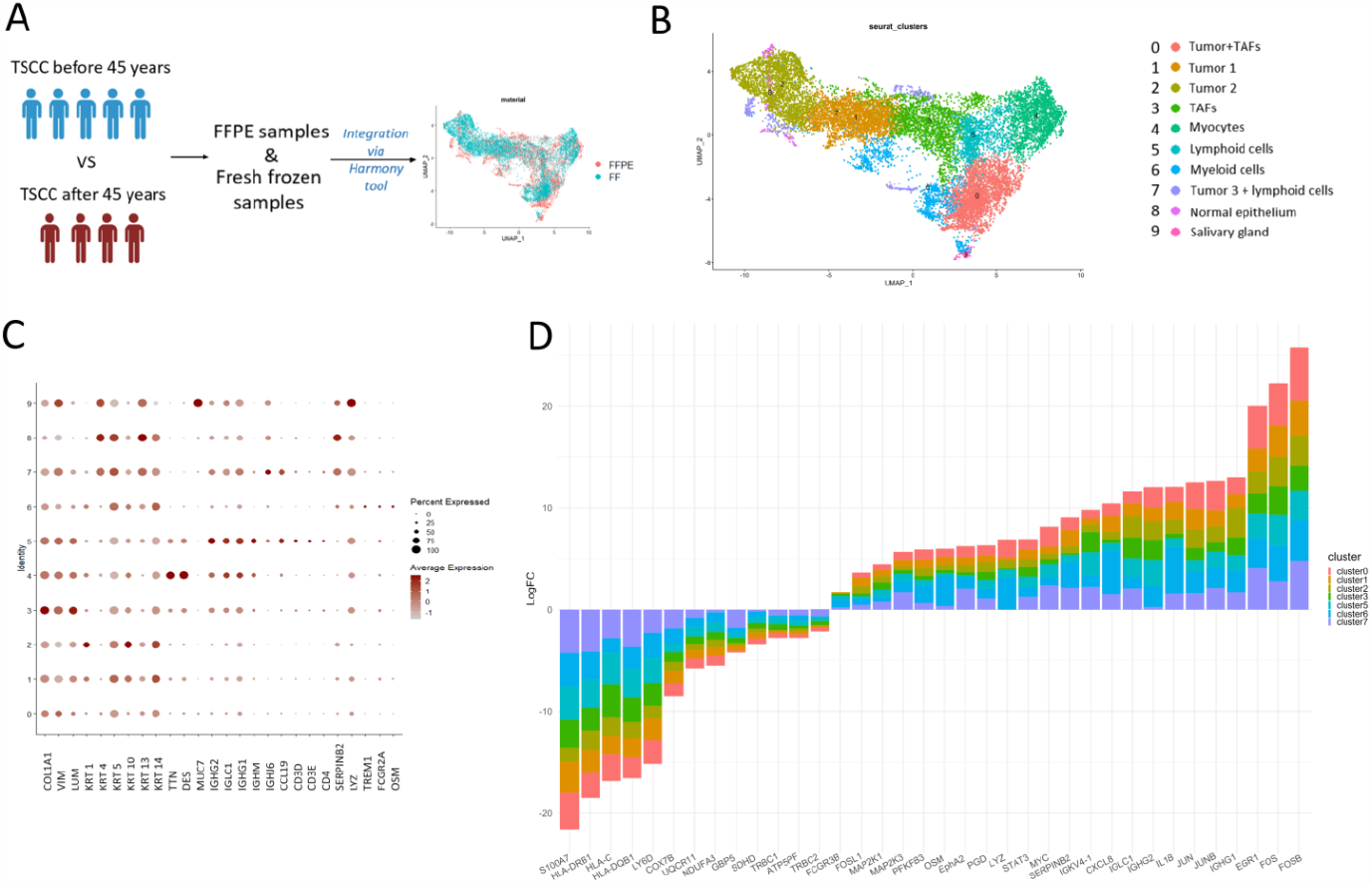
Spatial transcriptomics of early-onset TSCC. A, Sampling strategy, cohort overview, and sample integration; B, Overview of the identified cell types; C, Cluster identification by cell type markers; D, Consensus plot displaying the cumulative average logFC for genes (adj. p < 0.001) differentially expressed between early- and late-onset TSCCs.

We performed ST on 9 surgically resected TSCC samples from unique patients using the 10x Genomics Visium platform. Because each spot contains more than one cell, the accuracy of spatial transcriptomics is lower than that of single-cell RNA sequencing. We used the Harmony algorithm for dataset integration and found shared cellular types across tumors. A total of 9 cell clusters were partitioned into eight major cell types, including two tumor cell clusters, lymphoid and myeloid immune cells, TAFs, muscles, normal epithelium, and salivary gland cells (Figure 1B). Two tumor cell clusters were distinguished based on the expression of the *KRT1, KRT4, KRT5, KRT10, KRT13*, and *KRT14* genes (Figure 1C). Tumor microenvironment cells were detected as clusters of TAFs and lymphoid and myeloid immune cells (Figure 1B, 1C). Two clusters were mixed with fibroblasts (“tumor + TAFs”) and lymphoid cells (“tumor 3 + lymphoid cells”; Figure 1B, 1C). Next, we focused on tumor and immune cell types as key players in tumorigenesis.

### Transcriptome features of tumor cells in early-onset TSCC

As compared to elderly patients, TSCC cells of young adults were reliably distinguished by increased expression of mitogen activated kinase (MAPK) pathway genes: *EGR1, IL1B, MAP2K1, MAP2K3, FOS, FOSB, JUN, JUNB, FOSL1*, and *FOSL2* (Figure 1D). In contrast, the expression of oxidative phosphorylation genes (*ATF5PF, SDHD, GBP5, NDUFA3, UQCR11, COX7B*, and *S100A7*) was decreased in TSCC cells of young adults, while the glycolysis marker *PFKFB3* was increased (Figure 1D).

### Transcriptome features of immune cells in early-onset TSCC

Early-onset TSCC was also distinguished by increased expression of the *SERPINB2, LYZ*, and *OSM* genes corresponding to tumor-associated macrophages (TAMs) and the *FCGRB3* and *PFKFB3* genes being markers of myeloid-derived suppressor cells (MDSCs) (Figure 1D). In contrast, the T-cell markers (*LY6D, TRBC2*, and *TRBC1*) and the *HLA-DRB1, HLA-DQB1, HLA-C* genes associated with antigenic presentation were downregulated in early-onset TSCC, while the expression of immunoglobulin markers (*IGHG2, IGHG1, IGLC1*, and *IGKV4-1*) was upregulated (Figure 1D).

### Developmental trajectories of tumor cells in early-onset TSCC

Four cases with early-onset TSCC (excluding the FF sample) showed similar differentiation hierarchies with a progenitor cluster of cells (Figure 2). Progenitor cells were highly differentiated due to the expression of the *KRT16* and *KRTDAP* genes and expressed MAPK (*IL1B, TNFRSF21, IGF1R, IGFBP3, MAP2K3*, and *MAP2K1*) and tumor growth factor beta (*TGFB1*) pathways genes. The second and third clusters of 1 and 4 cases as well as the second clusters of 2 and 3 cases were decided to characterize together as an intermediate cluster owing to the similarity of gene signature expression. The intermediate cluster originated from progenitor cells was characterized by strong MAPK pathway activation and canonical TGFb signaling inhibition expressing the *DCN, FST*, and *THBS2* genes (inhibitors of the *TGFB1)* and the TGFb signaling activation through the *BMP2* gene. In addition, tumor cells of this cluster demonstrated features of the epithelial-endothelial transition and the activation of vascular mimicry through upregulation of the *EPHA2, FN1, MMP14*, and *COL1A1* genes and the JAK-STAT signaling pathway (*STAT3* and *MYC* genes). The last cluster showed low differentiation due to decreased expression of the *KRT16* and *KRTDAP* genes and retained expression of the MAPK pathway and vascular mimicry genes.

**Figure 2.**
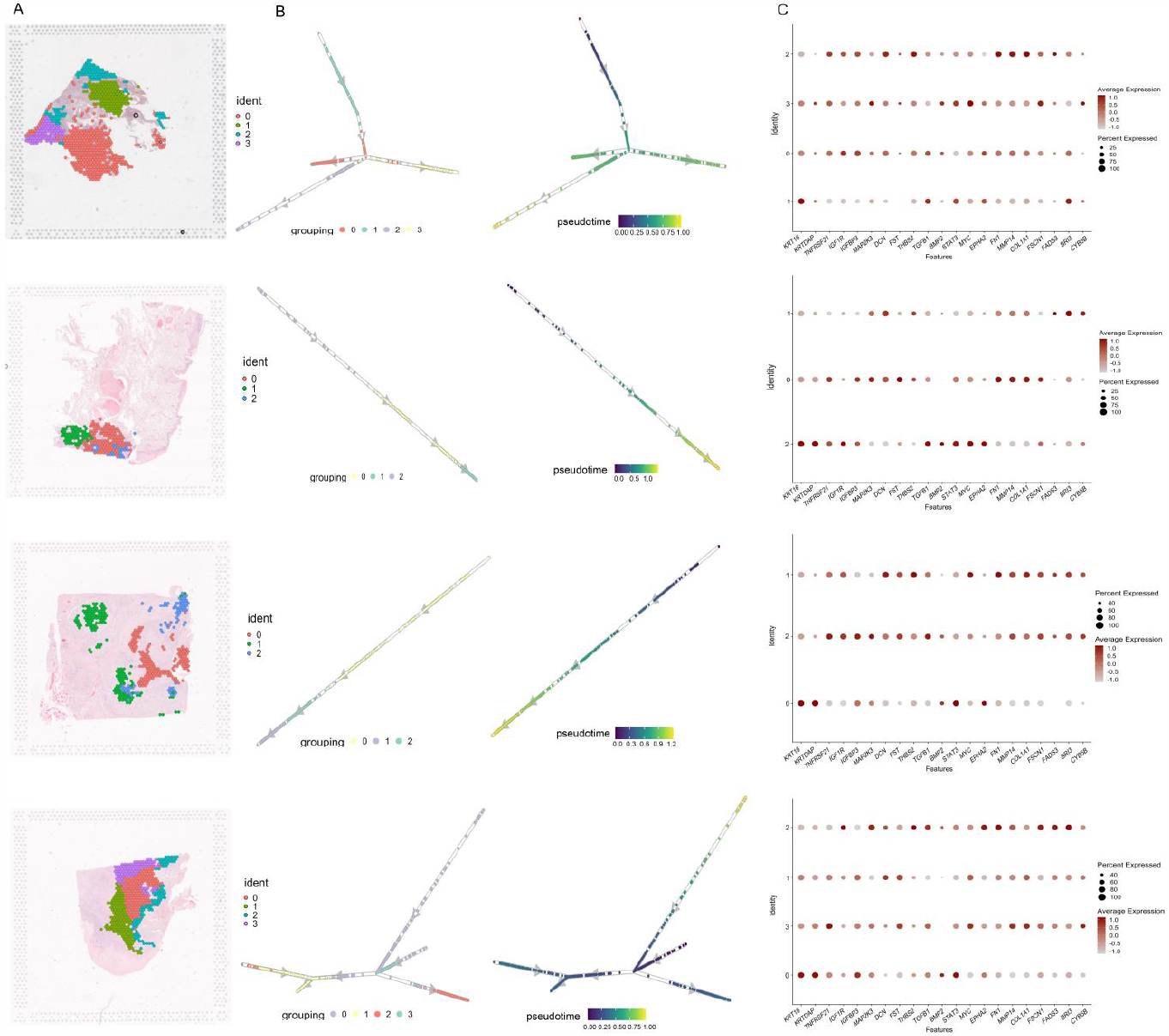
Tumor cell trajectories in early-onset TSCC and major driver genes. A, Visualization of tumor cell clusters in tissue sections; B, Directions of the cellular trajectories determined by unsupervised pseudo-time; C, Dotplot showing expression of top genes in each tumor cluster timepoint.

The genes specific to the last cluster of four early-onset TSCC cases were analyzed for association with overall survival using the Human Protein Atlas Kaplan-Meier plotter tool [12]. The focus on the last cluster was related to the suggestion that during the transition of tumor cells from a lower to higher malignant state, some genes are silenced, while others become active, thus predicting aggressiveness and early recurrence of cancer [13, 14]. High expression of the *FSCN1, FADS3, BRI3*, and *CYB5B* genes was found to be associated with worse overall survival of head and neck cancer patients (Supplementary Figure S1).

### Spatial transcriptome features of the invasive front in early-onset TSCC

In contrast to TSCC in elderly patients, the border between tumor and normal tissue in 4 of 5 young patients was infiltrated by cells enriched with the immunoglobulin genes such as *IGHG2, IGHG1, IGLC1, IGKV4-1*, and *JCHAIN* corresponding to plasma cells (Figure 3A). These cells were presented in tertiary lymphoid structures (TLSs), which were absent in the tumors of elderly patients. The TLS gene signature was also observed in early-onset TSCC (Figure 3B) and included the above-mentioned immunoglobulin genes and T-cell markers (*TRBC1, TRBC2*, and *IL7R*).

**Figure 3.**
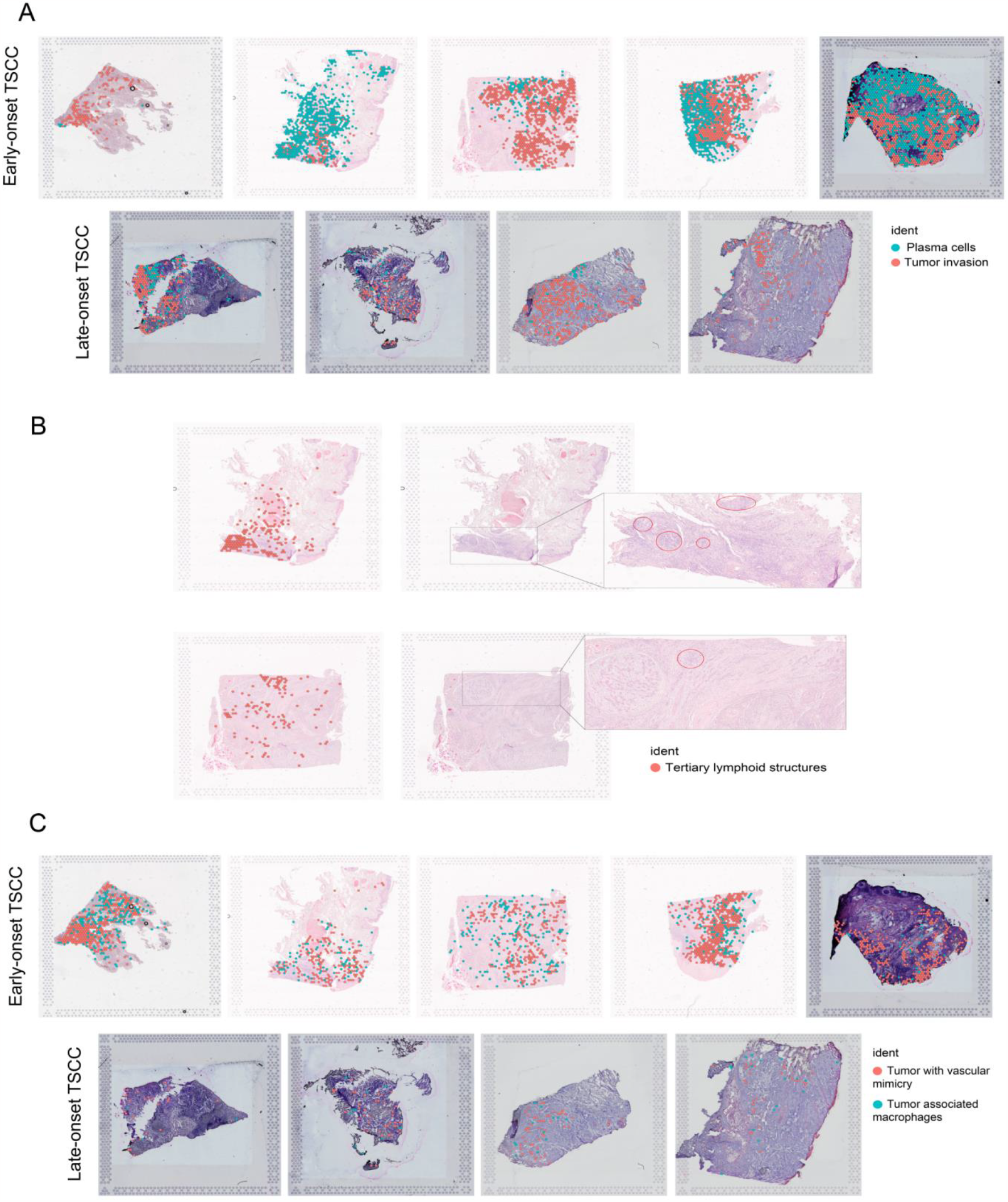
Spatial transcriptomic features of tumor and microenvironment regions in early- and late-onset TSCC. A, Spatial co-localization of tertiary lymphoid structures with gene signatures of plasma cells (*IGHG2, IGHG1, IGLC1, IGKV4-1*, and *JCHAIN*) and T lymphocytes (*TRBC2, TRBC1*, and *IL7R*); B, Plasma cell gene signature (*IGHG2, IGHG1, IGLC1, IGKV4-1*, and *JCHAIN*) at the border of tumor and normal tissue in early-onset TSCC; C. Markers of vascular mimicry (*ITGA5, ITGB1, MAP2K3, STAT3, MYC, EPHA2, FN1, MMP14*, and *COL1A1*) and TAMs (*CD68, SERPINB2*, and *OSM*) are co-located in solid tumor cell structures at the invasive front.

TSCCs of young patients were distinguished by an increased expression of the *ITGA5, ITGB1, MAP2K3, STAT3, MYC, EPHA2, FN1, MMP14*, and *COL1A1* genes corresponding to the vascular mimicry associated pathway (Figure 3C). This expression signature was found in solid tumor cell structures located on the border with normal tissue and corresponding to the areas of tumor invasion. Interestingly, regions of vascular mimicry were enriched with macrophage markers: *CD68, SERPINB2*, and *OSM*.

## DISCUSSION

The mechanisms of early-onset TSCC are unclear due to a lack of complex investigation of tumor and microenvironment cell compartments. This study used spatial transcriptome sequencing to characterize molecular features of tumor and microenvironment cells in early-onset TSCC and to reveal potential mechanisms of cancer development. Unlike elderly patients, TSCC of young adults was found to be enriched with immunosuppressive gene signature, oxidative stress, the MAPK and JAK-STAT molecular pathways, as well as with plasma cells, TAMs and vascular mimicry features at the invasive front (Figure 4).

**Figure 4.**
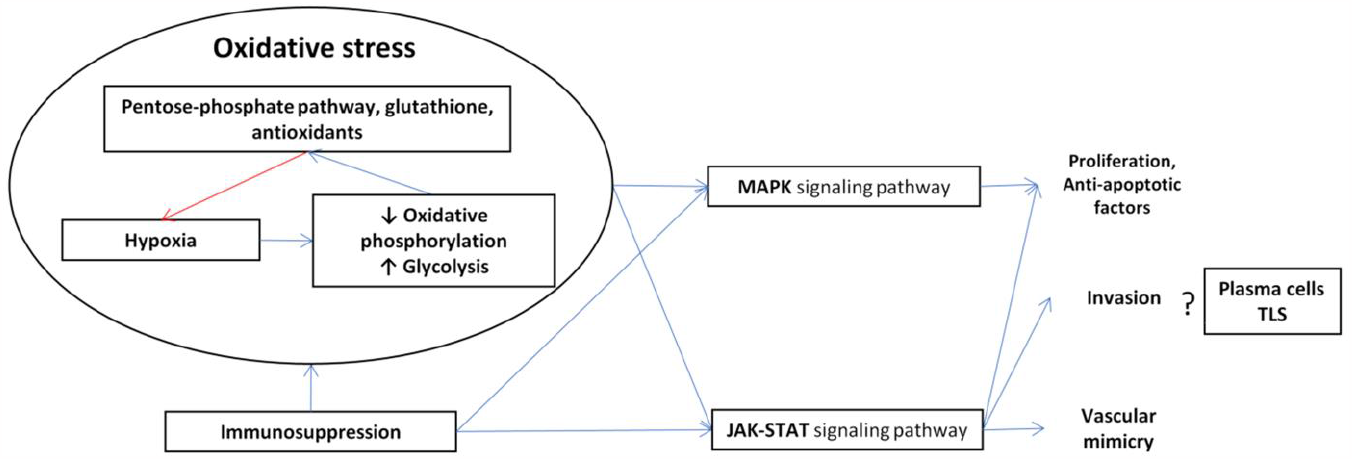
Hypothetical model of early-onset TSCC pathogenesis. Hypoxia and oxidative stress maintained by immunosuppression seem to be drivers of early-onset TSCC. Oxidative phosphorylation induces tumor development through the MAPK and JAK-STAT signaling pathways that contribute to tumor cell proliferation, apoptosis inhibition, invasion, and vascular mimicry. Plasma cells and TLSs are enriched in the invasive front; however, their role in early-onset TSCC pathogenesis is not yet clear. Red color refers to possible inhibition action; blue color – possible promotion action.

Early-onset TSCC demonstrates extremely downregulated oxidative phosphorylation and upregulated glycolysis. This may be due to the rapid growth of the tumor and its aggressiveness [15, 16]. The key molecular player in early-onset TSCC development is most likely the MAPK signaling pathway, which can be activated by immunosuppressive cells [17, 18] and upregulated in response to hypoxia and oxidative stress [15]. This finding is also supported by our previous analysis of the TCGA data for early-onset TSCC [9]. Activation of the MAPK signaling pathway is known to cause the transformation of normal cells into a tumor phenotype [19], and its targeting has the potential to arrest tumor growth. Currently, certain inhibitors are being used in clinical trials, and some of them have led to an improvement in the objective response rate, for example, in non-small cell lung cancer [20]. Non-canonical TGFb signaling pathway through the *BMP2* gene upregulation was found in early-onset TSCC; however, current data is scarce to speculate if this signaling is activated [21].

Tumor microenvironment is well known to contribute to tumorigenesis, cancer progression, and therapy resistance. Many cell types and functionally distinct cell populations comprise the tumor microenvironment demonstrating tumor-supportive or tumor-suppressive roles [22-24]. The early-onset TSCC microenvironment was found to be enriched in MDSCs possessing immunosuppressive and tumor-promoting properties. This cancer also demonstrated a decrease in the expression of the major histocompatibility complex (*HLA-DRB1, HLA-DQB1*, and *HLA-C*) and lymphocyte marker (*LY6L, TRBC2*, and *TRBC1*) genes characterizing an important link in the immune response and antigen presentation. The analysis of the TCGA data also showed PD-L1/PD-1-mediated immunosuppression and inhibition of phagocytosis in TSCC at young age [9]. In light of these findings, we suggest that early-onset TSCC have a pronounced immunosuppression. Moreover, our data calls into question the view that immune checkpoint inhibitors can be effective for treating young patients with TSCC.

The invasive front of the tumor represents an area where tumor cells actively contact with and invade into surrounding normal tissue contributing to further metastasis and recurrence. The invasive edge of early-onset TSCC was found to be enriched with vascular mimicry demonstrating upregulation of the JAK-STAT signaling pathway genes and surrounded by TAMs, which may induce or support this process [25-27]. The JAK-STAT signaling pathway was also found to be enriched in TSCC at young age according to the TCGA data analysis [9]. Vascular mimicry or blood vessels formation by tumor cells is often found in solid tumors contributing to tumor cell survival and metastasis [28]. In addition, the tumor border is enriched with plasma cells and lymphocytes aggregating to TLSs indicating a better adaptive immune response in young patients [29, 30]. Nevertheless, as said before, early-onset tongue tumor cores are characterized by an increased number of immunosuppressive cells. Further research is needed to reveal the mechanisms of this phenomenon and its importance for disease outcome.

In conclusion, the present study shows that TSCC in young adults has a distinct spatial transcriptomic features and potential molecular mechanisms of pathogenesis compared to older patients and should be identified as a separate clinical form with its own treatment, post-treatment follow-up tactics, and prognosis. Further studies should elucidate the contribution of MAPK and JAK-STAT signaling pathways and local immune defense characteristics to early-onset TSCC and the development of new diagnostic and prognostic markers and therapeutic targets.

## MATERIAL AND METHODS

### Tumor sample collection and annotation

The study included primary tumors from two cohorts of patients (n=9). The clinical data on patients are presented in Supplementary Table S1. The first cohort included 4 patients (aged < 45 years) with TSCC treated at the Cancer Research Institute of Tomsk National Research Medical Center of the Russian Academy of Sciences. The second cohort included 5 patients (1 patient aged < 45 years and 4 patients aged > 45 years), tumor samples of which were obtained from the Alberta Cancer Research Biobank. All patients did not receive neoadjuvant treatment. Tumor samples from the first cohort were formalin-fixed, paraffin-embedded (FFPE), and the second ones were fresh frozen (FF). The group of 5 patients aged up to 45 years (4 FFPE and 1 FF samples) was compared with the group of 4 patients aged over 45 years (4 FF samples) (Figure 1A).

The study was approved by the Ethics Committee of the Cancer Research Institute of Tomsk National Research Medical Center of the Russian Academy of Sciences and Health Research Ethics Board of Alberta – Cancer Committee and performed according to the guidelines of Declaration of Helsinki and the International Conference on Harmonization’s Good Clinical Practice Guidelines (ICH GCP). All subjects gave written informed consent in accordance with the Declaration of Helsinki.

### Spatial transcriptomic profiling

Visium spatial gene expression slides and reagents (Visium Human Transcriptome Probe Set v1.0, 10x Genomics, USA) for spatial transcriptomics assay were used according to manufacturer instructions. To assess the quality of the selected FFPE blocks, RNA was isolated by PureLink FFPE RNA Isolation Kit (Invitrogen, USA) from serial 10μm-thick tissue sections. Determination value index DV_200_ that represents the percentage of RNA fragments larger than 200 nucleotides with respect to all RNA fragments was calculated using the 4150 TapeStation system (Agilent, USA). Samples which had a DV_200_ ≥ 50% were considered good quality and selected to proceed with the experiment. Five-micrometer FFPE sections of TSCC samples were placed on a Visium Gene Expression slide (10x Genomics, USA). Deparaffinization, hematoxylin and eosin staining, and decrosslinking were performed according to the protocol of Visium Spatial Gene Expression for FFPE (10x Genomics, USA). Leica Aperio AT2 histoscanning station (Leica, Germany) was used to visualize tissue sections on slides. After incubation with the Probe Hybridization Mix (10x Genomics, USA), the tissues were permeabilized and captured. Gene Expression (GEX) libraries were generated for each section and then sequenced on a GenoLab M (GeneMind Biosciences, China). The Visium data on FF samples were kindly provided by Pinaki Bose and Rohit Arora [11].

### Spatial transcriptomics data processing

Sequencing reads for FF tissues were aligned using the 10x Genomics Space Ranger 1.3.1 pipeline (10x Genomics, USA) and 2.0.0. for FFPE tissues to the standard GRCh38 reference genome. Seurat package (v.4.3.0) was used for the following analysis. For each sample, the spatial file and H5 file obtained from SpaceRanger were loaded by the function “Load10X_spatial”. We filtered each Seurat Objects to exclude low-quality and outlier spots. The filtering parameters are presented in Supplementary Table S2. For spatial transcriptomics data obtained from FF samples, mitochondrial genes were excluded from the analysis. Data were normalized using the “NormalizeData” function with a scale factor of 10,000. Variable genes were identified using the “FindVariableFeatures” function. Data were scaled and centered using the “ScaleData” function. Harmony package version 0.1.1 was used to integrate data correcting by material type (FF or FFPE). For optimal results, genes not captured by FFPE probes were removed from the FF spatial transcriptomics data prior to integration. Cell clusters were identified via the “FindNeighbors” and “FindClusters” functions using a resolution of 0.5 for integrated object. “FindAllMarkers” function was used to find differentially expressed genes between all clusters and according to patient group. We used a Wilcoxon rank-sum test with a logFC > 0.5 and an adjusted P value < 0.001.

### Cell type annotation

Marker genes were used to determine whether clusters belonged to known cell types: tumor-associated fibroblasts (TAFs) -*COL1A1, VIM*, and *LUM*; tumor cells - *KRT1, KRT4, KRT5, KRT10, KRT 13*, and *KRT 1;* myocytes – *TTN* and *DES;* salivary glands cells – *MUC;* lymphoid cells - *IGHG2, IGHG1, IGLC1, IGHM, IGHJ6, CCL19, CD3D, CD3E*, and *CD4;* myeloid cells – *SERPINEB2, LYZ, TREM1, FCGR2A*, and *OSM* (Figure 1C). The “DotPlot” function (Seurat package) was used to visualize the expression of marker genes across different clusters (Figure 1C).

### Trajectory inference

After isolating tumor cell-containing spots from young patient samples, each sample was independently used for maturation trajectory inference using dynverse packages, which provide more than 50 different methods for trajectory inference. Normalized tumor cell data and UMAP coordinates were used as input. We also entered prior information about start cell identifier, which was chosen based on the expression of the genes responsible for epithelial cell differentiation (*KRTDAP, MKI67, CD44*, and *S100A14*), as well as information about the *KRTDAP* gene that is known to be important in oral cell differentiation [31]. The most suitable and rapid method for our dataset determined by applying “guidelines” function was the PAGA Tree. The visualization of the trajectories and pseudotime as a 2D graph structure was done with the “plot_graph” function (dynplot package). A standard “SpatialDimPlot” function (Seurat package) was used to visualize clusters of tumor cells on slides.

### Statistical analysis

The Human Protein Atlas Kaplan-Meier plotter tool was used to evaluate the correlation between the gene expression and the survival rates in head and neck cancer [32].

## Acknowledgments

This work was supported by the Russian Science Foundation (# 22-15-00308).

---

**Figure S1.**
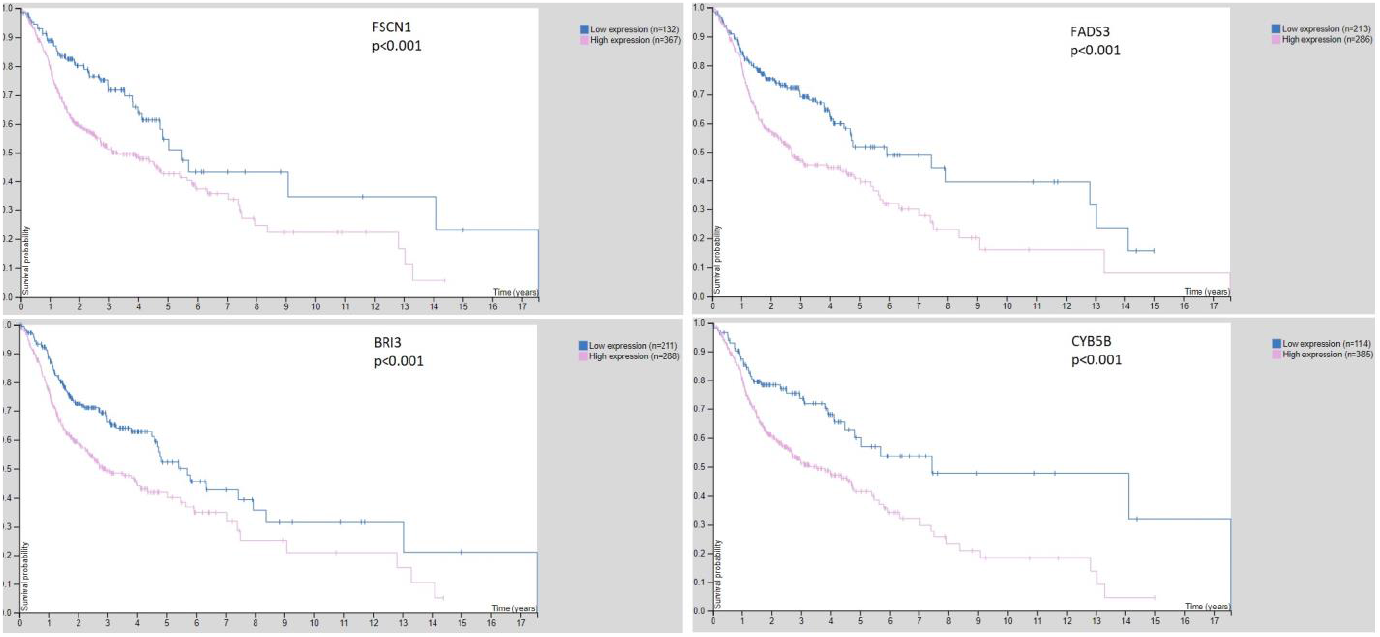
Correlation between the *FSCN1, FADS3, BRI3*, and *CYB5B* gene expression and overall survival in patients with head and neck cancer patients.

**Supplementary Table 1. Patient clinical information for all samples used for analysis**

**Supplementary Table 2. The filtering parameters**

