## Supplementary Table 1. Patient clinical information for all samples used for analysis for "Revealing Molecular Mechanisms of Early-Onset Tongue Cancer by Spatial Transcriptomics" (from another version of this preprint: version 1, 2023-10-16)

| Patient ID | Center of sequencing | Sex | Age | Smoker Status (Years Smoked) | T stage | N tage | M stage | Differentiation | HPV status |
|---|---|---|---|---|---|---|---|---|---|
| Outs 1 | Tomsk National Research Medical Center, Russia | Female | 31 | Never Smoked | 3 | 1 | 0 | mod diff | negative |
| Outs 2 | Tomsk National Research Medical Center, Russia | Female | 33 | Never Smoked | 2 | 1 | 0 | well diff | negative |
| Outs 3 | Tomsk National Research Medical Center, Russia | Male | 37 | Never Smoked | 3 | 1 | 0 | well diff | negative |
| Outs 4 | Tomsk National Research Medical Center, Russia | Female | 43 | Never Smoked | 3 | 0 | 0 | well diff | negative |
| Outs 5 | University of Calgary, Canada | Male | 53 | Current Smoker | 2 | 2 | 0 | mod diff | negative |
| Outs 6 | University of Calgary, Canada | Male | 50 | Ex Smoker | 1 | 0 | 0 | well diff | negative |
| Outs 7 | University of Calgary, Canada | Male | 37 | Never Smoked | 2 | 2 | 0 | well diff | negative |
| Outs 8 | University of Calgary, Canada | Male | 61 | Never Smoked | 2 | 0 | 0 | mod diff | negative |
| Outs 9 | University of Calgary, Canada | Female | 49 | Never Smoked | 2 | 0 | 0 | mod diff | negative |
