## Supplementary Table 2. The filtering parameters for "Revealing Molecular Mechanisms of Early-Onset Tongue Cancer by Spatial Transcriptomics" (from another version of this preprint: version 1, 2023-10-16)

| Sample | Sample group | nFeature_Spatial | nCount_Spatial | percent.mt |
|---|---|---|---|---|
| Outs 1 | young | 200 -6000 | 500-20000 | -* |
| Outs 2 | young | 200 -6000 | 500-40000 | -* |
| Outs 3 | young | 200 -6000 | 500-20000 | -* |
| Outs 4 | young | 200 -7000 | 500-25000 | -* |
| Outs 5 | old | 200-7500 | 500-35000 | < 10 |
| Outs 6 | old | 200-8000 | 500-50000 | < 10 |
| Outs 7 | young | 200-9000 | 500-50000 | < 10 |
| Outs 8 | old | 200-6000 | 500-30000 | < 10 |
| Outs 9 | old | 200 -5000 | 500-20000 | < 6 |

Note: * FFPE probes do not capture mitochondrial genes
